# The Pesticide Chlorpyrifos Increases the Risk of Parkinson’s Disease

**DOI:** 10.1101/2025.03.13.643184

**Authors:** Kazi Md Mahmudul Hasan, Lisa M Barnhill, Kimberly C Paul, Chao Peng, William Zeiger, Beate Ritz, Marisol Arellano, Michael Ajnassian, Shujing Zhang, Aye Theint Theint, Gazmend Elezi, Hilli Weinberger, Julian P Whitelegge, Qing Bai, Sharon Li, Edward Burton, Jeff M Bronstein

## Abstract

**Background and Purpose:** Pesticides have been associated with an increased risk of Parkinson’s disease (PD), but it is unclear which specific pesticides contribute to this association and whether it is causal. Since chlorpyrifos (CPF) exposure has been implicated as a risk factor for PD, we investigated its association to incident PD and if this association is biologically plausible using human, rodent, and zebrafish (ZF) studies.

**Methods:** The association of CPF with PD was assessed using the UCLA PEG study (829 PD and 824 control subjects), and proximity-based exposure estimates from living or working near agricultural CPF use. For the mammalian studies, 6 months old male C57BL/6 mice were divided into two groups, CPF and controls, for open field, rotarod, and wire hang behavioral testing. Mice were then exposed to CPF in an inhalation chamber (0.65-2.9 mg/m^3^/day) for 6 hrs./day 5 days/wk., whereas control mice were exposed to vehicle alone. Behavioral tests were performed before and 2.5 months after CPF exposure following a 3-day washout. Mice were then perfused for immunohistochemical analysis. For the mechanistic studies, ZF embryos were treated with CPF (250 nM) 24 hours post fertilization for 5-7 days. Behavioral testing was performed using the Viewpoint Imaging System. Neuronal loss and microglial activation were determined using immunohistochemistry. Neuronal autophagic flux was determined using autophagy modulators in GFP-LC3 transgenic ZF and Western blots.

**Results:** Long-term residential CPF exposure was linked to an increased risk of developing PD with an odds ratio of 2.68 (CI 1.58-4.55). Mice exposed to aerosolized CPF developed motor impairment and a significant loss of dopaminergic neurons in the substantia nigra and activation of microglia. TH positive neurons in the substantia nigra (SN) had significantly higher levels of phosphoserine 129 (pS129) α-synuclein (α-syn), a marker for pathological phosphorylated α-syn, and ubiquitin. In contrast, neither pS129 α-syn or ubiquitin accumulated in TH neurons in the VTA after CPF exposure. Consistent with the mice data, CPF exposure resulted in impairment of locomotor activity and selective loss of aminergic neurons in ZF. We also found an increase in neuronal apoptosis and microglial activation. Importantly, dopamine neuron loss was found to be at least partially dependent on γ1-synuclein (closest functional homologue to human α-syn) as neuronal loss did not occur in γ1-synuclein knockout ZF. Using an *in vivo* ZF assay, we found impaired autophagic flux and an increase in lysosomal labelling within the zebrafish brain. CPF exposure also led to elevated γ1-synuclein and p62 (autophagic cargo protein) levels consistent with impaired degradation. Furthermore, induction of autophagy was protective, supporting the hypothesis that impaired autophagic flux is at least partially responsible for neuron loss following CPF exposure.

**Conclusions:** CPF exposure is associated with an increased risk of developing PD and this association is likely causal since PD-like pathology was recapitulated in animal models. Furthermore, impaired autophagic flux appears to underly this toxicity, a pathway implicated in the pathogenesis of PD.

## Introduction

Parkinson’s disease (PD) is a slowly progressive neurodegenerative disorder manifested by motor dysfunction and cognitive decline. The main pathological hallmarks of PD are the selective loss of dopaminergic (DA) neurons in the substantia nigra pars compacta (SNpc), the development of fibrillar cytoplasmic inclusions, also known as Lewy bodies and Lewy neuritis, and inflammation. The major component of Lewy bodies and neurites is misfolded, highly ubiquitinated and phosphorylated α-synuclein (α-syn). The etiology of PD appears complex and involves both genetic and environmental interactions. A minority of PD cases are caused by mutations in several genes including *SNCA*, *LRRK2*, *GBA*, *VPS35*, *RAB32*, and *PINK1* that code for proteins involved in proteostasis, mitochondrial function, and inflammation.^1–3^ The etiology of the majority of PD cases is not known but likely involves environmental toxins. Pesticide exposure as a class of toxins is one of the strongest and best documented risk factors associated with an increased risk of PD but few individual chemicals have been identified that confer this increased risk.^4, 5^ It is essential to identify specific chemicals in order to determine if the associations are causative and what are the mechanisms of toxicity.

Several studies have investigated the association of environmental toxins with the development of PD but there are challenges in determining if an association is causing the altered risk. It is now clear that PD develops over decades, so any exposure assessment should be performed prior to when the pathology starts. Once a toxin has been identified to be associated with altered risk, it is then necessary to perform animal experiments to determine the biological plausibility that this exposure is initiating and/or propagating disease pathology. These studies should test whether exposures in animals can recapitulate some of the clinical and pathological features of PD and determine the mechanisms by how they act. For example, rotenone is one of very few environmental toxins that have met all these criteria. Agricultural exposure to rotenone is associated with an increased risk of developing PD and rats injected with rotenone develop motor abnormalities, α-syn neuronal inclusions, DA neuronal loss, and inflammation.^6, 7^ Furthermore, rotenone is known to act by inhibiting mitochondrial complex I.^6, 7^

Chlorpyrifos (CPF; O,O-diethyl O-3,5,6-trichloro-2-pyridyl phosphorothioate), is a broad spectrum and extensively used organophosphate pesticide in agriculture to control insect pests. Its target as a pesticide is through the inhibition of acetylcholinesterase which leads to neurological dysfunction and death of the insects. CPF is generally applied to crops by spraying and therefore human exposure is primarily through inhalation for agricultural workers. Although CPF had been extensively used as a pesticide in the United States, few studies have investigated its association with the risk of developing PD. Furthermore, the effects of CPF exposure in animals have been inconsistent, and no clear mechanism of toxicity has been established.

Here we report that exposure to CPF is associated with an increased risk of developing PD in a large community-based case-control study. Mice exposed to CPF using a novel inhalation method that recapitulates human exposures, caused impaired motor behavior, loss of DA neurons, increased pathological α-syn, and inflammation. Using transgenic ZF, we found that CPF was neurotoxic by disrupting autophagic flux and that DA neuron toxicity was dependent on γ1-synuclein (closest functional homologue to human α-synuclein). Together, these studies strongly implicate that exposure to CPF as a risk factor for developing PD and modulators of autophagy are a promising therapeutic target.

### CPF exposure and its association with PD

To assess the association of CPF with PD risk, we utilized the Parkinson’s Environment and Genes (PEG) study (n = 829 PD patients; n = 824 controls). As shown in Table 1, PD patients were on average slightly older than controls, a higher proportion were men, had European ancestry, and were never smokers. We estimated ambient exposure due to living or working next to agricultural facilities applying CPF over a 30+ year period. We observed positive associations between CPF and PD with exposure estimated at residential and workplace and over different exposure time windows (Fig. 1D, Table 1). The strongest association was with the longest duration at the workplace with an OR of 2.74 (CI 1.55, 4.89). Importantly, CPF exposures over the 20 to 10 years prior to disease onset were mor strongly associated with PD than the 10-year period prior to PD onset, consistent with the theory that PD pathology begins decades prior to clinical symptoms are appreciated.

**Figure 1:**
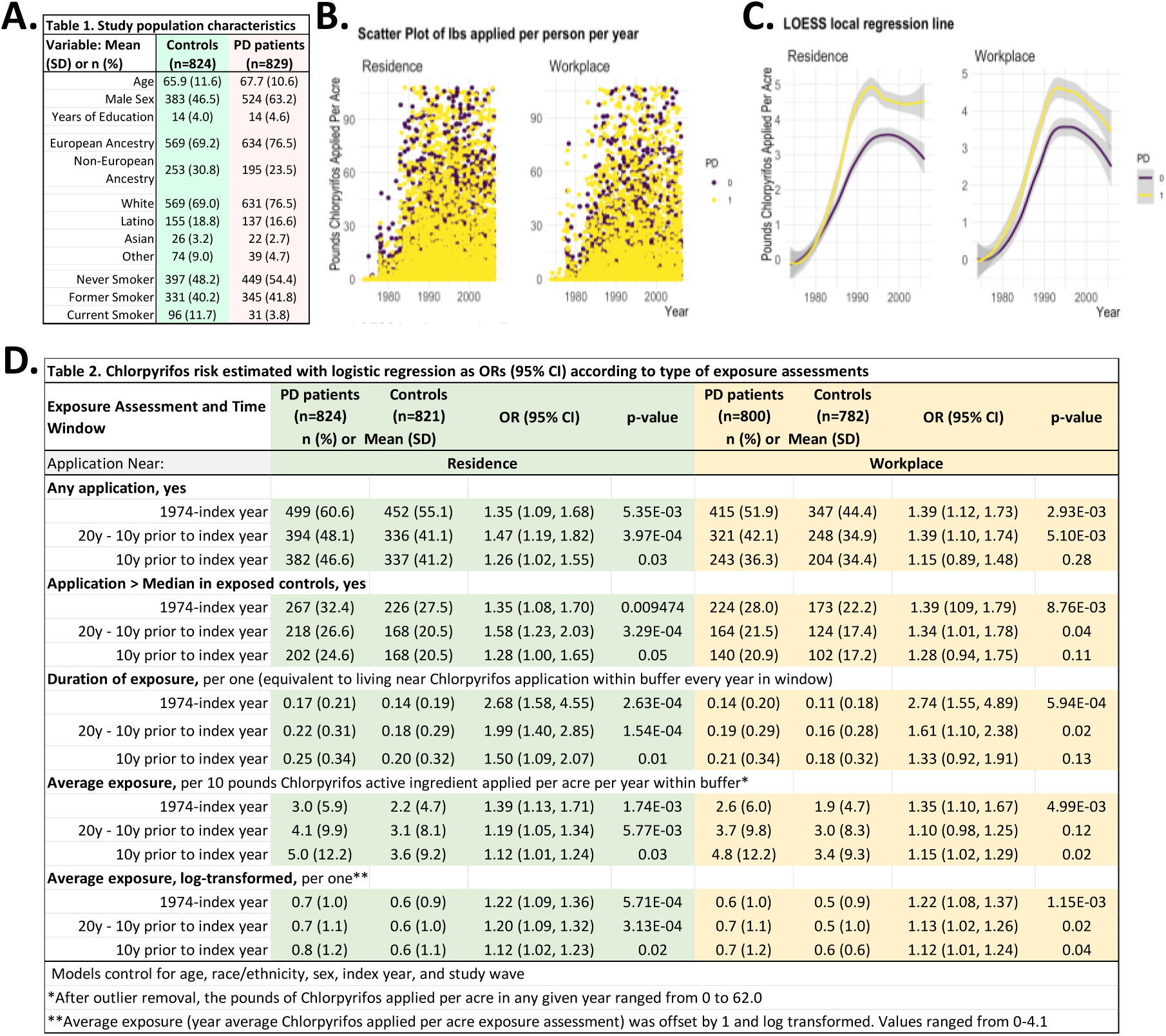
Pounds of agriculturally applied CPF active ingredient per acre, per subject 1974–2007. **(A)** Demographic characteristics of PD patients and non-PD participants **(B)** Scatter plot of pounds active ingredient applied per acre for each participant each year within 500 m of residential address and workplace address. **(C)** Smoothed trend line from loess local regression based on data shown in plots. Lb, pounds; LOESS, locally estimated scatterplot smoothing. **(D)** CPF risk estimated with logistic regression as ORs (95% CI) according to type of exposure assessments.

### Effects of inhaled CPF on motor behavior in mice

The majority of human pesticide exposure is primarily through inhalation which escapes the 1^st^ pass circulation to the liver as with oral ingestion and therefore reduces its metabolism. In order to model human exposures, mice were exposed to aerosolized CPF or vehicle alone in closed chambers five days a week for 11 weeks. We found in preliminary experiments that female mice were much more resistant to CPF than males and therefore, only male mice were tested in the current study (data not shown). The concentrations of CPF used were determined empirically (highest dose that was well tolerated) and it was increased over time as the mice adapted to the exposures. CPF-exposed mice maintained their body weight as did the ethanol vehicle controls until the last week of the experiment (Fig 2). The concentration of CPF in the chamber reached 650 µg/m3/day initially and peaked at 2900 µg/m3/day by week 11 (Fig. 2B). Concentration of CPF in the brains of exposed mice were measured by liquid chromatography-tandem mass spectrometry and reached 1.44 to 2.42 ng/mg tissue (Fig 2C).

**Figure 2:**
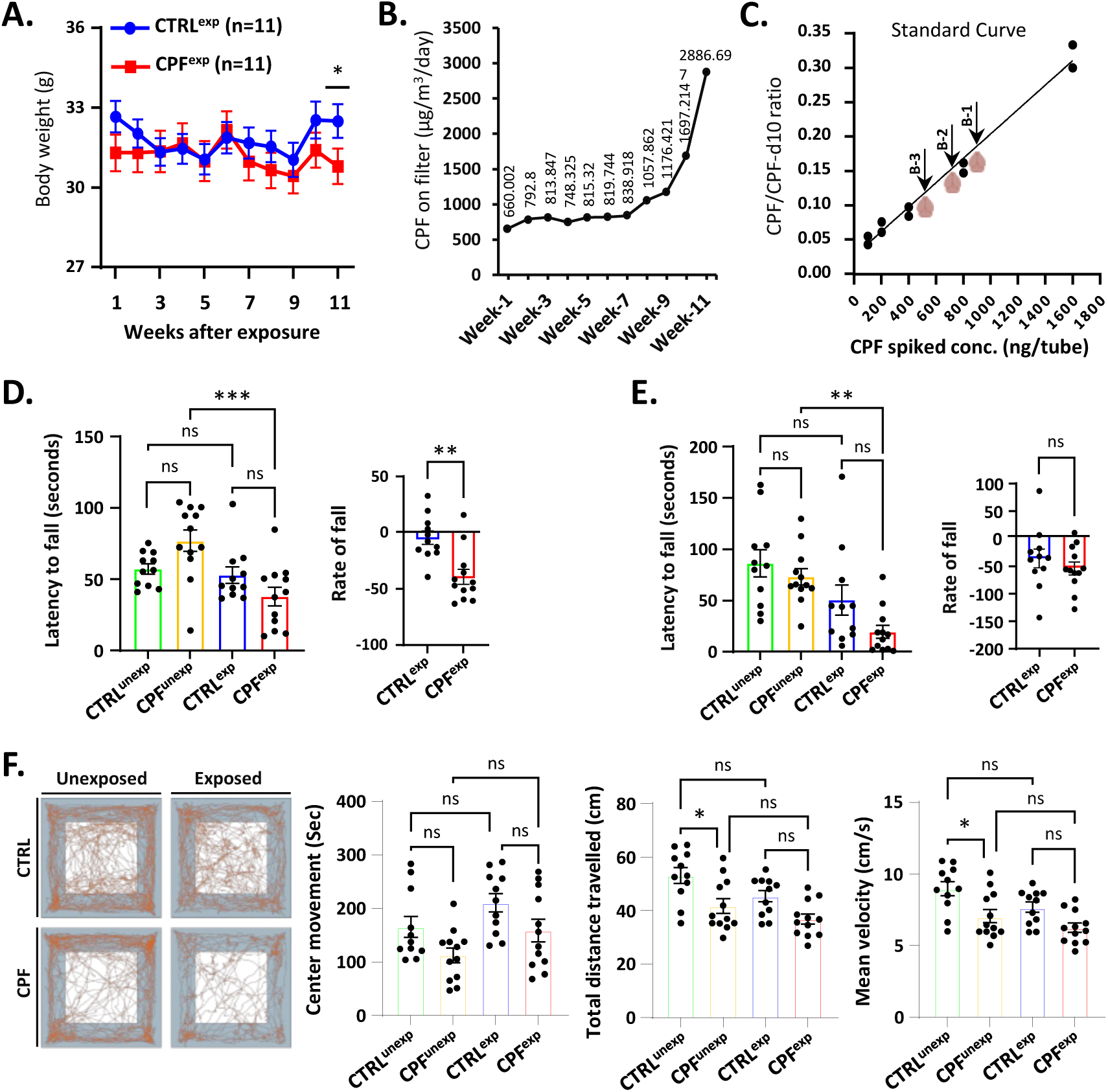
CPF exposure parameters and behavior assay on C57BL/6J mice. (A) Mice body weight recorded before and after CPF exposure. (B) CPF concentrations deposited in the exposure chamber. (C) CPF standard curve and mouse brain concentrations (arrows) (D) Accelerating rotarod performance, with average latency to fall (left) and falling rate (right) of control and CPF mice before and after exposure. (E) Wire hang test performance, with latency to fall (left) and falling rate (right) of control and CPF mice before and after exposure. (F) Open field performance. From left to right: Representative trajectories of individual control and CPF mice before and after exposure, mean center time, total distance traveled, and mean velocity. *p < 0.05, **p < 0.01, ***p < 0.001, and ns=not significant. P-values were calculated using the two-way analysis of variance (ANOVA). Control group (CTRL): n=11; Experimental group (CPF): n=12.

Mouse behavior was analyzed before the initiation of CPF exposures and 3 days following the last day of exposure using rotarod, wire hanging and open field testing. The 3-day washout was used to ensure that the behavior was not altered by CPF or vehicle and likely reflected underlying pathology. Both the exposed and control mice performed worse after 11 weeks presumably due to aging or possibly the stress of the exposures. CPF mice deteriorated more than controls in all 3 tests although only the rotarod test reached statistical significance (Fig 2D-F).

### CPF exposure leads to loss of DA neurons

A core pathological feature of PD is the relatively selective loss of DA neurons which results in impaired motor behavior. We determined the total number of DA neurons in control and CPF exposed mice using stereological methods. The number of DA neurons in the CPF exposed brains in the SNpc decreased by 26% ± 10.1 compared to the control animals but were unchanged in the VTA (Fig. 3A). Phosphorylated α-syn at serine 129 (pS129) is a biomarker for pathological forms of α-syn and is elevated in PD brains. We found elevated levels of pS129 staining and ubiquitin in DA neurons of the SNpc compared to controls whereas staining was similar in VTA neurons (Fig 3 C-F). Western blot analysis confirmed that pS129 was elevated 1.5-fold in the detergent soluble of the CPF brains compared to controls (Fig 3). Staining for TH in the striatum was reduced in the CPF exposed brains relative to controls consistent with DA neuron loss in the SNpc but the difference did not reach statistical significance (Fig. 3G).

**Figure 3:**
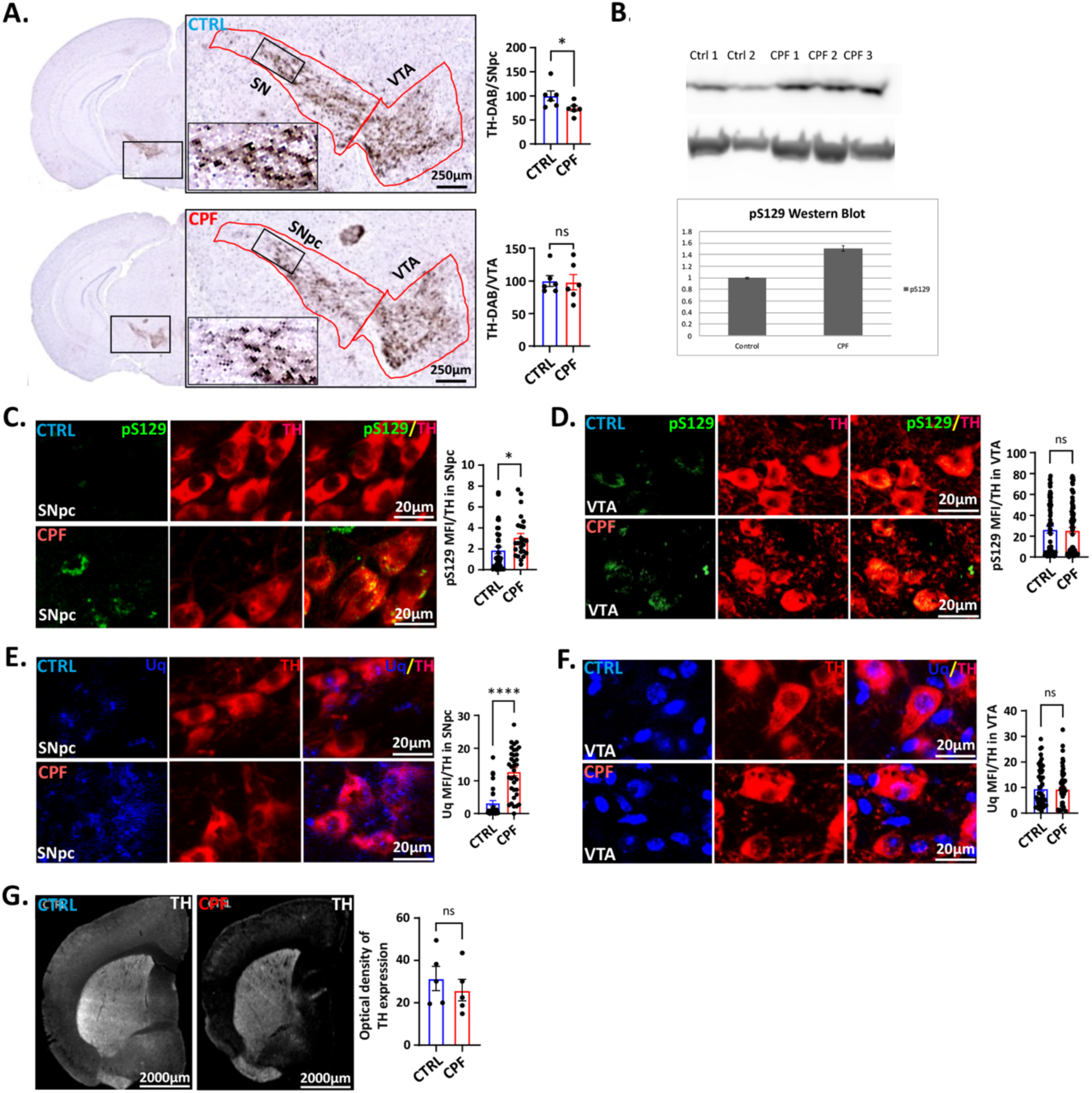
Effects of CPF inhalation on mice DA neurons. (A) Immunohistochemical staining and histogram of the number of TH-positive cells in the SNpc and ventral tegmental area (VTA) of mice brain. Scale = 250 μm. CTRL: n=5; CPF: n=6. (B) α-Syn pS129 was increased in CPF soluble fractions. (C, D) Representative SNpc, and VTA images with histogram of pS129 α-syn positive aggregates in CPF-exposed mice. Scale = 20 μm. (E, F) Representative SNpc, and VTA images with histogram of ubiquitin staining. Scale = 20 μm. (G) Representative TH-stained striatum. Scale = 2000 μm. Statistical significances are represented by standard error of mean (SEM) with asterisks: *p < 0.05, ****p < 0.0001, and ns=not significant. P-values were calculated using nonparametric Mann–Whitney t test. CTRL: n=5; CPF: n=6.

### Microglial take on an activated morphology after CPF exposure

CNS inflammation is a common feature in PD and may contribute to its pathogenesis.^8, 9^ Microglia change their shape and become more rounded in response to various stimuli such as environmental toxins and infections. ^9–12^ To investigate whether microglial take on an activated morphology after CPF exposure, we determined the mean size, and perimeter of Iba1-stained cells using the method of Fernandez-Arjona et al.^13^ We found that CPF-exposed microglia were more rounded and had shorter projections in the SNpc and VTA but not in the SNpr (Fig 4). These morphological changes are consistent with activated microglia similar to that seen PD brains.

**Figure 4:**
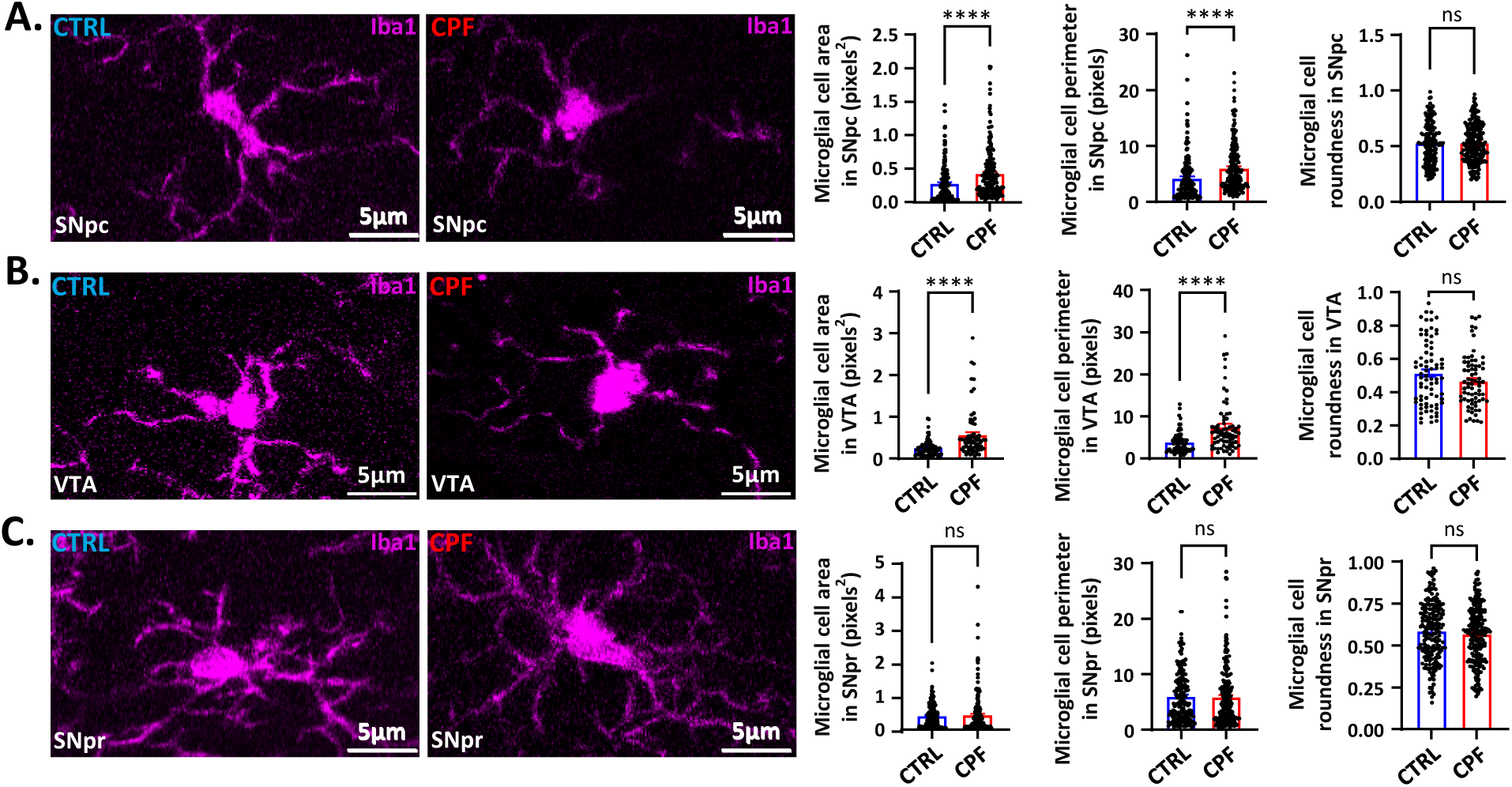
CPF inhalation and neuroinflammation in mice brain. (A-C) Representative images of 40× Z-stacked photomicrograph of Iba1 staining used for microglia analyses at the SNpc, VTA, and SNpr. Images were converted to binary with randomly selected microglia. Histogram represented the microglial size (area, and perimeter), and roundness. Scale bar = 50 mm. Statistical analysis performed by two-tailed, non-parametric Mann-Whitney U-test. Statistical significance is represented by standard error of mean (SEM) with asterisks: ****p < 0.0001, and ns=not significant. CTRL: n=5; CPF: n=6.

### CPF exposure induces selective neurotoxicity to DA neurons in zebrafish

We have found that exposure to CPF is associated with an increased risk of developing PD and that mice exposed to CPF through inhalation develop many PD features including motor dysfunction, loss of DA neurons, α-syn pathology and inflammation. To determine the mechanism of CPF neurotoxicity, we utilized transgenic ZF that are transparent and contain a well-developed DA system in the larval stage. Dechorionated ZF, 24 hours post fertilization (hpf), were exposed to 0 to 100µM CPF for 6 days to determine its overall toxicity and found that all fish died at 30µM or higher by day 7 (Supplementary Fig. 2A and B). To determine whether CPF is toxic to aminergic neurons including DA neurons, we have utilized the transgenic (Tg) *vmat2*:GFP line that expresses green fluorescent protein (GFP) under the control of vesicular monoamine transporter 2 (*vmat2*) promoter to monitor aminergic (DA, noradrenergic, and serotonergic) neuronal integrity as previously described.^14–16^ We used the lowest concentration at which we observed decreased locomotor activity (250 nM) for all subsequent experiments. A stereotypical pattern of swimming response to light was observed in vehicle-treated embryos and CPF-treated fish at 5 dpf but the CPF-exposed ZF swam slower than controls in the dark at 7 dpf (Supplementary Fig. 1C, D). This pattern of ZF behavior is consistent with DA neuron loss^17, 18^.

The integrity of the aminergic system in ZF was determined at 5 and 7 dpf after CPF (250 nM) exposure. The number of aminergic neurons in the telencephalic (Tc), diencephalic (Dc), and TH positive neurons in the Dc clusters at 5 and 7 dpf ZF embryos was significantly reduced (Fig. 5A-C, Supplementary Fig. 3A, B). Interestingly, increased apoptosis was observed in the telencephalic and diencephalic regions of 5 dpf embryos but not in the larvae at 7 dpf (Supplementary Fig. 3A and B, 4A and B) suggesting that this is the time of active neuronal death. Since PD brains show some selectivity for DA neuron loss, we tested the selectivity of loss after exposure to CPF. Unlike *vmat2* neurons, Rohon-Beard neurons in 7 dpf Tg(*isl1*[ss]:Gal4-VP16,UAS:eGFP)^zf154^ larvae expressing GFP in their tail regions showed no significant changes after CPF treatment (Fig. 5D). Thus, CPF exposure led to the selective loss of aminergic neurons similar to that seen in PD.

**Figure 5:**
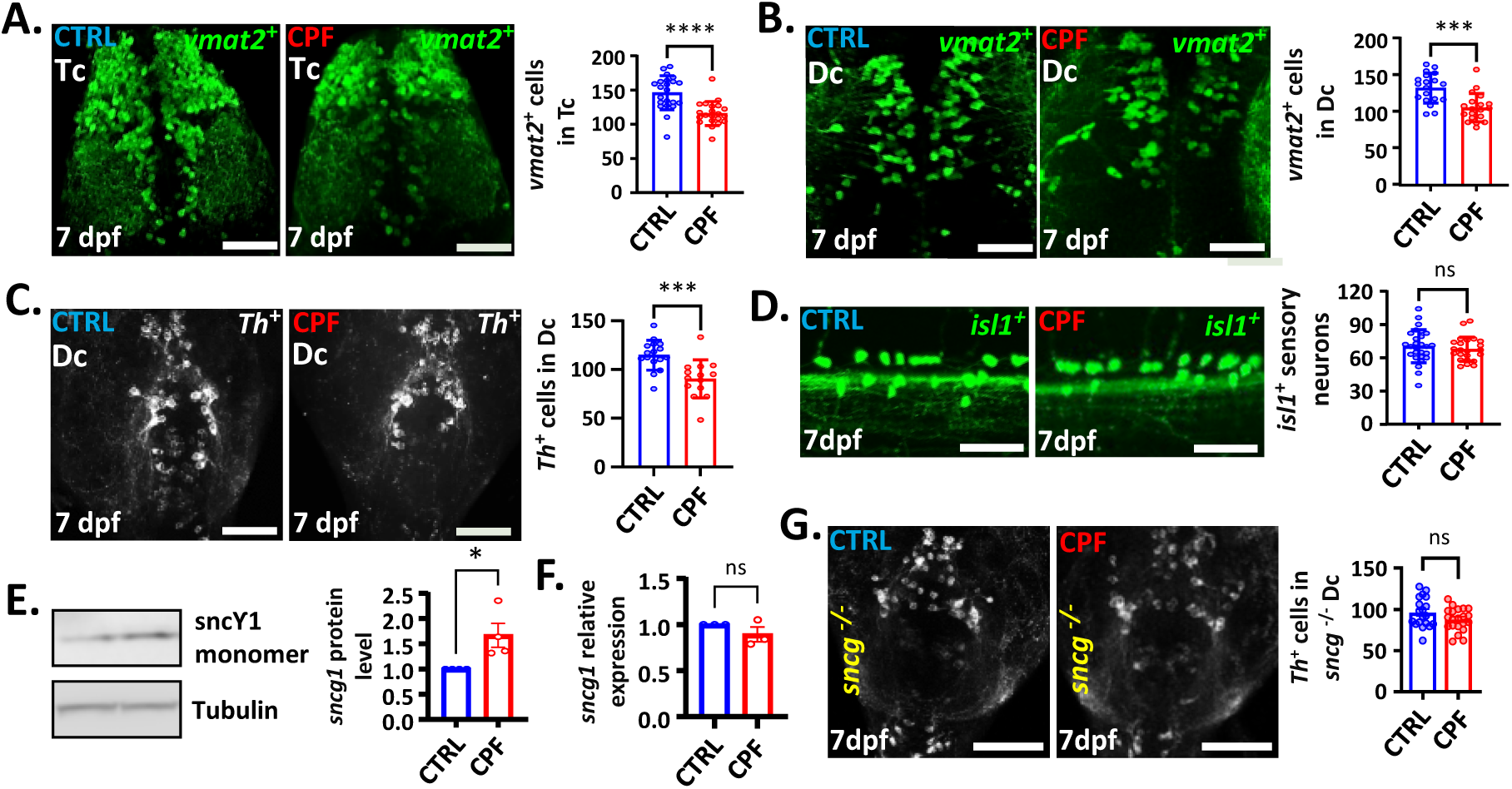
CPF induces selective loss of DA neurons in ZF embryos. CPF exposure leads to reduced *vmat2*:gfp positive neurons (A, B) and TH positive neurons (D) in the telencephalon (Tc), and diencephalon (Dc) (dorsal view). CPF Did not cause loss of non-aminergic isl1+ neurons (D; lateral view). Scale bar = 50 μm. E) CPF treatment resulted in an increase in γ1-syn determined by western blot while γ1-syn mRNA levels were unchanged (F). (G) Representative Th immunostaining showed that sncγ1 knockout zebrafish are protected against CPF-induced DA neuron loss. Scale bar = 50 μm. Statistical analysis performed by two-tailed, non-parametric Mann-Whitney U-test. Statistical significances are represented by standard error of mean (SEM) with asterisks: *** = p < 0.001, **** = p < 0.0001, ns = not significant.

### Exposure to CPF activates microglia but does not contribute to neurotoxicity

Another pathological feature of PD is CNS inflammation which we also found in the CPF-exposed mice. Since we can easily reduce the number of microglia in ZF using morpholinos, we tested if microglia contribute to CPF-induce neuron loss. ZF larvae exposed to CPF showed higher number of microglia that had a more rounded cell body and shorter processes at 7 dpf compared to controls (Fig. 6A). Furthermore, the maximum branch length, the number of branches, and the number of junctions of microglia decreased. These structural changes in microglia are indicative of a more activated state much like what we saw in the mice. It has been reported that activated microglia have more and/or larger lysosomes.^19^ Using live imaging and lysotracker green in *mpeg1*-mCherry ZF, we found that the fluorescence intensity (MFI) of lysotracker labeling that co-localized with microglia was significantly increased following exposure to CPF (Fig. 6B). Collectively, CPF exposure resulted in structural and functional changes in larval ZF brains consistent with microglia activation.

**Figure 6:**
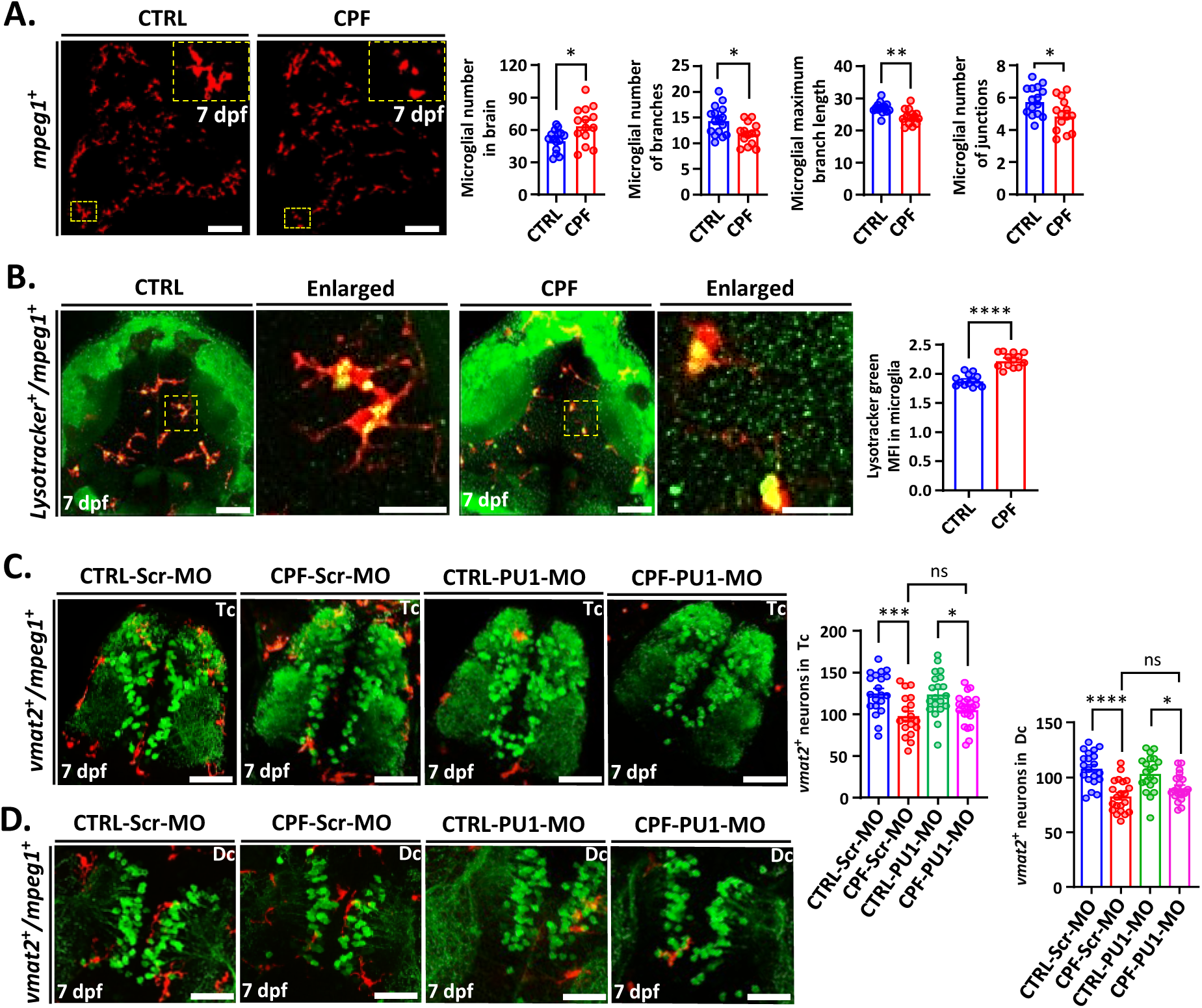
Microglia are not required for CPF neurotoxicity in ZF. (A) Tg(*mpeg1*:mcherry) zebrafish embryos were treated with 0.25 µM CPF at 24 hpf for 6 days. CPF exposure significantly increased the number of microglia in the brain, while the microglial maximum branch length, number of branches and number of junctions were significantly decreased at 7 dpf. Scale bars: 75 μm. (dorsal view). (B) CPF-exposed *mpeg1*:mCherry ZF brains have higher lysotracker staining in the brain as a whole and in microglia. Scale bars: 25 μm, and 5 μm (Enlarged). Statistical analysis performed by two-tailed, non-parametric Mann-Whitney U-test. (C, D) Injection of PU1 morpholino (MO) blocked microglial development *mpeg1*:mCherry embryos compared to scrambled MO-injected embryos in both Tc and Dc. PU1 and scrambled morpholino-injected fish showed no significant difference in CPF-induced DA neuron loss in Tc and Dc. Scale bar = 50 μm. Statistical analysis was performed by two-way ANOVA with Sidak’s multiple comparisons test. Statistical significances are represented by standard error of mean (SEM) with asterisks: * = p < 0.05, ** = p < 0.01, *** = p < 0.001, **** = p < 0.0001, ns = not significant.

To evaluate whether microglia are involved in CPF-induced neurotoxicity, we inhibited microglial development using PU1 morpholino (MO) before CPF exposure. Neither scrambled control or PU1 MO injections caused significant toxicity, nor did injection significantly affect the number of surviving neurons in the embryos. As shown in Figures 6C and 6D, injection of PU1 MO resulted in marked reductions (87%-91%) of *mpeg1*:mcherry-positive microglia through 7 dpf in both Tc and Dc regions of the brain. Reduction of microglia had no significant effect on CPF neurotoxicity although there was a non-significant trend for the microglia deficient ZF to have less of a loss than ZF with microglia. Aminergic neurons were reduced by 22% in the telencephalon with no injection, 21% loss with control morpholino (scrambled) injection, and 14% loss with PU1 injection following CPF treatment (Fig. 6C). Similar results were found in the diencephalon (Fig. 6D). These results suggest that microglia do not contribute significantly to CPF neurotoxicity at least in this acute model.

### γ1-Synuclein is necessary for CPF toxicity

Synucleins are neuronal proteins that comprise α-, β-, and γ-synucleins in mammals and α-syn aggregation plays a central role in PD pathophysiology.^20^ ZF do not contain an ortholog of human α-syn although ZF γ1-synuclein (γ1-syn) has a similar function as α-syn.^21^ In order to determine if γ1-syn was involved in CPF toxicity in ZF, we exposed larvae to CPF for 6 days and found that its levels were increased compared to controls (Fig. 5E). We then utilized γ1-syn knockout ZF to determine if this increase in γ1-syn contributed to DA neuron loss. Interestingly, CPF exposure did not result in DA neuron loss in γ1-syn knockout ZF (Fig. 5G). Thus, γ1-syn was required for CPF neurotoxicity in ZF.

### Reduced autophagic flux contributes to CPF neurotoxicity

CPF exposure led to an increase in γ1-syn levels that was due to either an increase in expression or decreased degradation. We measured its expression using qRT-PCR after CPF exposure and found a non-significant trend for decreased γ1-syn mRNA compared to controls (Fig. 5F). These results suggest that γ1-syn degradation is impaired in CPF-exposed ZF. α-Syn is degraded by both the ubiquitin proteasome system (UPS) and autophagy. Since aggregated α-syn is primarily degraded by autophagy, we focused on the effects of CPF on this system.^22^ Here, we utilized the Tg(*elavl3*:eGFP:*map1lc3b*) transgenic line to measure autophagic flux in neurons *in vivo* after CPF exposure using published methods with minor modifications.^23^

A significant reduction in GFP-positive autophagosome (*Lc3-II*) puncta was observed after treatment with CPF (Fig. 7A). We then determined the autophagic flux by determining the number of autophasomes after 1 hr. treatment with saturating concentrations of bafilomycin A1and we found that CPF significantly reduced flux (Fig. 7A). A reduction in flux by CPF exposure was further supported by an increase in p62 protein levels, while p62 expression was unchanged (Fig. 7B and C). If disruption of autophagy contributed to DA injury, stimulating autophagy should be neuroprotective. We utilized calpeptin to induce autophagic flux (Fig. 7A and Supplementary Fig. 4), and we found that it rescued the neuron loss caused by CPF (Fig. 7D-F). This was confirmed both by counting *vmat2*-GFP neurons and *Th* immunohistochemistry (Fig. 7D-F). Thus, CPF exposure led to reduced autophagic flux that contributes to DA neuron loss.

**Figure 7:**
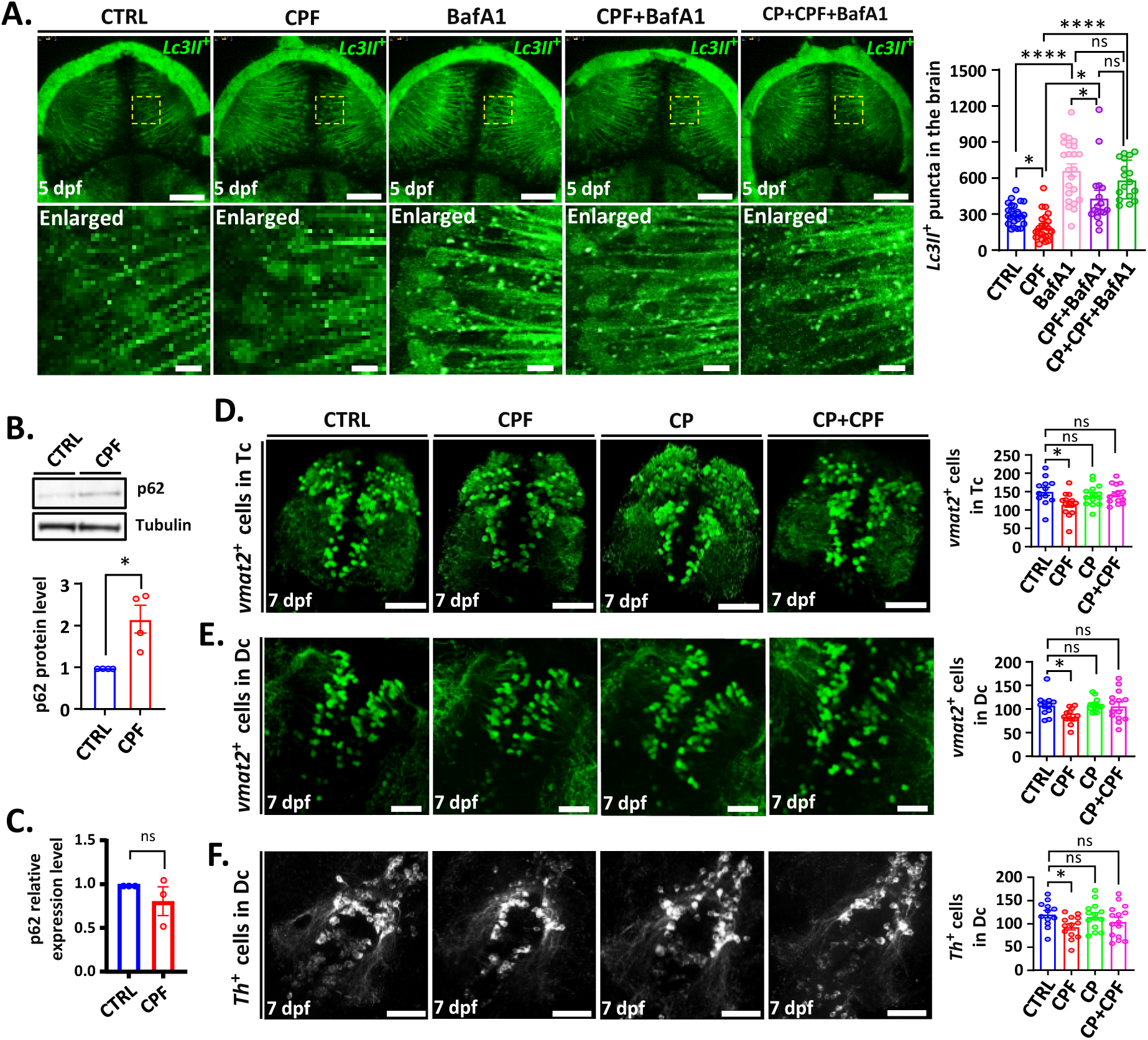
CPF inhibits autophagic flux that leads to DA neuron loss in ZF. The number of *Lc3II-*GFP positive puncta in the midbrain were counted in 5 dpf larvae with and without Baf1A. (A). Statistical was performed by two-way ANOVA with Brown-Forsythe and Welch’s multiple comparisons test. Scale bar = 50 μm, and 5 μm (Enlarged). (B) p62 levels increased in CPF-exposed zebrafish embryos. (C) Quantitative PCR showed reduced p62 expression in CPF treated embryos. (D-F) *Vmat2*-GFP positive neurons after CPF-exposure with and without calpeptin treatment to induce autophagy. Scale bar = 50 μm. Statistical analysis performed by two-tailed, non-parametric Mann-Whitney U-test for B and C. Statistical analysis showed the quantification of *vmat2*-GFP positive neurons using Brown-Forsythe and Welch’s multiple comparison ANOVA test. Statistical significances are represented by standard error of mean (SEM) with asterisks: * = p < 0.05, **** = p < 0.0001, ns = not significant.

## Discussion

The goal of this study was to determine if CPF exposure causes an increased risk of developing PD. We found that long-term exposure to CPF is associated with up to 270% increased risk of PD and that behavioral and pathological features of PD can be recapitulated in mice. Furthermore, we found that the mechanism of CPF toxicity includes decreased autophagic flux and accumulation of synuclein, both required for CPF-induced DA neuron loss. Taken together, these findings strongly support a causal association of CPF and increased risk of PD.

There are several strengths to this study which includes the PEG study. Exposures were estimated by using the Pesticide Use Report database in California, which provides detailed data on agricultural pesticide use since 1972. By knowing how much, where, and when CPF was sprayed, and where people lived and worked at the time of the spraying, we could estimate a subjects’ exposure. This approach has been well validated and is routinely used for air pollution research.^24^ The lag time between exposure and the diagnosis is extremely important since it is believed that the α-syn pathology begins decades prior to the core symptoms of PD become apparent. The association of CPF exposure with PD was stronger when we assessed exposures in the 10-20 yrs prior to the diagnosis than 0-10 years and strongest when the duration of the exposure were the longest (Table 2). Importantly, exposure assessment did not rely on a subjects’ recall which is often inaccurate and subject to recall bias. Another strength of the PEG cohort is that the PD diagnosis was confirmed by a movement disorders specialist who examined each subject, often multiple times, over several years. A weakness of our approach is that CPF exposure did not occur in isolation and despite controlling age, gender, and smoking, co-exposures with other pesticides or other unknown risk factors could not be accounted for. Our findings are consistent with another case-controlled trial with similar increases in risk found after CPF exposure.^25^

We next set out to determine if there is biological plausibility for the association of CPF with increased risk of developing PD using 2 different animal models. The effects of CPF on rodents have been mixed with some studies reporting a loss of DA neurons following oral and IP exposures.^26–28^ No animal study has tested the primary means by which humans are exposed, inhalation. Inhalation of toxins avoids the portal blood flow system to the liver that occurs with oral ingestion and therefore is not metabolized as rapidly. We exposed mice to inhaled CPF and found that it induced behavioral deficits, DA neuron loss, pathological α-syn, and inflammation, all characteristics of PD. The lesions in the DA pathway were only partial explaining why the behavioral changes and reduced TH staining were not profound. Mice were exposed for only 11 weeks which might explain this partial lesioning as compared to human exposures which occur over decades. We also tested middle-aged and not aged mice which are likely more vulnerable. Preliminary studies suggested that younger female mice are more resistant to CPF than older males which is why we report only on males in this study. Excess in male vulnerability to CPF is also consistent with the gender bias seen in humans.

We also found increased p-αsyn in DA neurons in the CPF mice relative to controls that was confirmed by Western blot. Although p-αsyn was increased and aggregates were present in some neurons, the state of α-syn did not appear to be primarily fibular. Syn 506, an antibody that reacts with fibrils, did not significantly stain the CPF-exposed brains (data not shown). The lack of more complex α-syn aggregates might have occurred for a number of reasons. More time might be necessary for them to form and/or microglia may have successfully been able to clear then.

An important component of establishing biological plausibility is determine the mechanisms by which a toxin acts. We utilized ZF for this purpose since they are transparent, easily manipulated genetically, and contain a well-developed DA system even in the larval stage. We found that CPF exposure at very low concentrations (250 nM) caused selective toxicity to DA neurons. This toxicity was at least partially due to γ1-syn since its levels were increased and DA neurons were unaffected when CPF was tested in γ1-syn knockout fish. The elevated levels were the result of decreased degradation since expression was unchanged. We found that CPF exposure resulted in a reduced autophagic flux that contributed to CPF-induced DA neuron loss. Interestingly, microglial activation did not appear to significantly contribute to DA neuron loss since markedly reducing their development with morpholinos had no effect on CPF neurotoxicity. It is very possible that microglia may contribute to pathology in more chronic models and in humans.

There are some weaknesses to the ZF model. They are developing at the time of exposure when PD is a disease of the aged. We do not think this invalidates our results since younger animals generally are more resistant to toxins relative to older animals suggesting that we may be underestimating CPF toxicity, not overestimating it. Another consideration is the difficulty in determining what a relevant concentration is and duration of exposure since those in humans clearly cannot be recapitulated. Exposures in the ZF studies were more acute than in those in the mice and the concentration was higher than those that we measured in the mouse brains. Since the primary purpose of using the ZF was to determine the mechanism of toxicity, we believe these weaknesses do not alter the validity of our findings.

## Conclusions

Exposure to CPF, a widely used pesticide, is associated with an increased risk of developing PD. Mice exposed to CPF, in a similar manner as humans, develop motor deficits and pathological characteristics of PD. We propose that the mechanism of CPF’s neurotoxicity is by reduced autophagic flux that results in increased αsyn. Taken together, CPF increases the risk of developing PD and its use should be restricted.

## Methods

### PEG study population

To assess CPF and PD associations, we used the Parkinson’s Environment and Genes (PEG) study (*n* = 829 PD patients; *n* = 824 controls). PEG is a population-based case-control study conducted in three agricultural counties in central California (Kern, Fresno and Tulare).^19^ Participants were recruited in two waves: Wave 1 (PEG1): 2000–07, *n* = 357 patients, *n* = 400 population-based controls; and Wave 2 (PEG2): 2009–15, *n* = 472 patients, *n* = 424 population-based controls. Patients were enrolled early in disease course [mean PD duration at baseline, 3.0 years (SD = 2.6)] and all were seen by University of California, Los Angeles (UCLA) movement disorder specialists for in-person neurological examinations, many on multiple occasions, and confirmed as having probable idiopathic PD.^20,21^ As shown in Table 1, PD patients were on average slightly older than controls and a higher proportion were men, had European ancestry, and were never smokers.

### CPF exposure assessment

We estimated ambient exposure due to living or working near agricultural CPF application, using pesticide use report (PUR) pesticide application data within a geographical information system (GIS)-based model.^22^ Since 1972, California law mandates the recording of commercial pesticide use in a database maintained by the California Department of Pesticide Regulation (CA-DPR) that includes all commercial agricultural pesticide use by pest control operators and all restricted pesticide use until 1989, and afterwards (1990–current) all commercial agricultural pesticide use. This database records the location of applications, which can be linked to the Public Land Survey System (PLSS), and the poundage, type of crop, and acreage a pesticide has been applied on, as well as the method and date of application. We combined the PUR with maps of land use and crop cover, providing a digital representation of historical land use, to determine pesticide applications at specific agricultural sites.^29^ PEG participants provided lifetime residential and workplace address information, which we geocoded in a multi-step process.^30^ We determined the pounds of CPF applied per acre within a 500-m buffer of the latitude and longitude representing each residential and workplace address per year since 1974, weighting the total poundage by the proportion of acreage treated (lbs/acre). For our study participants, this resulted in 12,904 annual records for residential and 8,968 for occupational site CPF exposure. After we identified and removed several extreme outliers (values >99th percentile of the distribution), the resulting data ranged from 0 to 108.44 lbs/acre.

### Experimental Animals

#### Mice

Six months old 24 male wild-type mice on C57BL/6 background weighting 29–33 g were purchased from the Jackson Laboratory, Sacramento, California, USA. The mice were housed in a vivarium with 12/12 hours of light and dark cycles. University of California, Los Angeles (UCLA) Animal Research Committee approved all experiments using animals in accordance with the US National Institutes of Health guidelines.

#### Chlorpyrifos (CPF) exposure by inhalation

Sixteen 26-week-old male mice were exposed to aerosolized CPF (Sigma #45395) and another 16 were exposed to room air passed through control vehicle containing 2% ethanol in identical chambers using custom-built whole-body chamber systems (CH Technologies, Westwood, NJ). The chambers hold up to 20 mice and is connected a Blaustein Jet Atomizer which produces aerosols that were tightly controlled with Lab Flow C Control box. Aerosols were generated from solutions containing CPF (0.2 mg/mL) in 40% ETOH at a flow rate into the atomizer of 15 mL/hr and aerosol generator air flow was 1.5 L/min and dilution air flow was 4.5 L/min. The concentration of CPF initially in the chamber was 76.63 ug/m^3^ and was gradually increased to 300 ug/m^3^. Mice were exposed 5 days/wk for 4 hrs/day for 11 weeks with total exposures of 0.65-2.9 mg/m^3^/day.

#### Behavioral testing

Behavioral testing was performed prior to CPF exposures and 3 days after the final day of exposure to washout any residual effects of the pesticide and vehicle.

#### Rotarod

Rotarod testing was carried out as previously described (PMID: 23161999). Mice were trained to run on the rotarod (Bioseb) with four 5-minute trials at 20 rotations per minute (rpm) separated by five minutes. One hour later, mice were placed on the rotarod at 4 (rpm) and the speed was linearly accelerated to 40 rpm over 300 seconds. The latency to fall and speed at which the mice fell was recorded. The mean latency to fall was calculated from two consecutive trials in each mouse.

#### Wire Hang Test

Wire hang testing was carried out according to a modified protocol as previously described.^31^ Briefly, the mice were placed on the top of a wire mesh (wire diameter 0.047” with 0.45” openings) connected to a stepper motor controlled by an Arduino microcontroller. At the start of the trial, the wire mesh was rotated ∼10 degrees in both directions, causing the mice to grip the wires, and then flipped 180 degrees so mice were suspended upside down. Latency to fall was measured using an infrared sensor placed below the wire mesh. If mice did not fall within 10 minutes, the trial was terminated. The average latency to fall was calculated from two trials performed 15 min apart.

#### Open Field

Open field testing was performed using a custom made acrylic square arena (20″ x 20″ x 15″) with a white floor that was evenly illuminated to 95 lux^32^. During the trial, the mice were allowed to explore freely for 10 minutes while videotaped. A deep neural network was trained on behavior videos to identify arena boundaries and anatomical markers on mice (nose, ears, and tailbase) using DeepLabCut (PMID: 31227823). The trained model was then used to automatically extract marker positions during each video frame. Mouse position was calculated as the centroid of the polygon between the mouse ear and tailbase markers, and custom written code in MATLAB was used to calculate mean velocity, distance traveled, and time in the center of the arena (defined as a square 20% of the arena diameter away from each wall).

#### Mouse Immunohistochemistry

After perfusion and fixation of 6 CPF and control mice, brains were embedded in paraffin blocks, cut into 6 mm sections and mounted on glass slides as previously described with some modifications ^33^. Briefly, sample containing slides were deparaffinized and rehydrated, followed by immersion in 88% formic acid for 5 minutes to enhance antigen detection. Then the sections were treated with 5% hydrogen peroxide (Sigma#H1009) in methanol (Fisher Scientific#A454-4) for 30 min to quench endogenous peroxidases. Then blocked in 0.1 M Tris with 2% fetal bovine serum (Tris/FBS) for 5 min. Both TH primary and secondary antibodies were diluted in Tris/FBS. Samples were then incubated with primary antibody (Abcam#AB76442) overnight at 4°C. Following washing, sections were incubated for 1 hour with species-specific biotinylated secondary antibody (VectorLabs#BA-9010) and then for 1 hour with VECTASTAIN® Elite® ABC-HRP Kits (VectorLabs#PK-6100). After rinsing with 0.1M Tris, the slides were incubated for 4 minutes with DAB reagent (VectorLabs#SK-4105). In subsequent steps, slides were rinsed and washed, counterstained in Hematoxylin (Fisher Scientific#6765001), then dehydrated and cover slipped. Slides were scanned into digital format on a Lamina scanner (Perkin Elmer) at 20x magnification. Digitized slides were then used for quantitative pathology. All TH positive neurons in the midbrain (SNpc and VTA) were counted blindly.

In a separate group of 6 pairs of mice, brains were quickly removed after perfusion with PBS and euthanasia and cut in half sagittal. One half of the brains were quickly frozen for biochemical studies and the other half were fixed in 4% paraformaldehyde for at least 24 hrs. Following dehydration in 30% sucrose, embedded in OCT (Sakura Finetek#4583) and 40 μM sections were collected. The sections were blocked with 3% FBS and 3% bovine serum albumin (Sigma#B-4287) and incubated overnight at 4^0^C in primary antibodies, washed with PBS, incubated in secondary antibodies overnight at 4^0^C before cover slipping. Primary antibodies including TH (Abcam#AB76442; 1:500 dilution), pS129 (Abcam#AB51253; 1:1000 dilution), ubiquitin (Santa Cruz Biotechnology#sc-271289; 1:500 dilution), and Iba1 (Synaptic Systems#234 308 Gp311H9; 1:500 dilution). The sections were imaged using a Zeiss Z.1 Light dual sided illumination sheet fluorescence microscope using ZEN software. Quantification of psyn was performed blindly by identifying TH neurons in the red channel and measuring fluorescence in the green channel.

### Microglia analysis in mice

Microglial analysis was conducted as previously described.^34^ Briefly, Images from 40x Z-stacks were analyzed in ImageJ and then binarized for microglial analysis. Using ImageJ FracLac plugin with hull and circle results, the outlines of individual cells were analyzed. A total of two “results” files were generated, which were tabulated from Excell: the “Box count summary” file displayed lacunarity and fractal dimension, whereas the “Hull and circle results” file displayed other morphological characteristics.

### Biochemical fractionation and immunoblot

Extractions were performed using a modified procedure of Henderson et al.^35^ Briefly, mouse half brains homogenized in 9 volumes high salt (HS) 1% Triton X buffer with protease and phosphatase inhibitors using Dounce homogenizers for 20 strokes. The samples were centrifuged at 100,000 x g for 30 min and supernatants were stored at -80. The pellets were homogenized in 9 volumes HS-Triton X buffer with 30% sucrose, centrifuged at 100,000 x g for 30 min at 4^0^C. Supernatants were again stored at -80^0^C. The pellets were homogenized in 9 volumes 1% Sarkosyl HS buffer, shaken for 30 min at room temperature, and spun at 100,000 x g for 30 min. The resultant pellet was washed in 1.4 mL PBS without Ca++ or Mg++, resuspended in DPBS, and sonicated. An equal volume of supernatants from each extraction were combined and run on a 12% Bolt Bis Tris gel (Thermo Fisher Scientific#NP0321BOX) for 50min at 150volts, transferred to a nitrocellulose membrane, and blocked in 5% non-fat milk. Primary antibodies for pS129 (Abcam#AB51253; 1:2000 dilution) and tubulin (Abcam#AB51253; 1:2000 dilution) were diluted in 1% non-fat milk with secondary donkey anti-rabbit horseradish peroxidase (HRP) (Santa Cruz Biotechnology#sc-2004; 1:2500 dilution) were diluted in 1% non-fat milk. Blots were developed in ECL Plus substrate for 5 minutes before imaging on Azure Biosystems imager and band intensity was quantified using ImageJ using the Analyze Gels tool. The pSer129 blots were developed with SuperSignal Atto substrate.

### Zebrafish husbandry, strains and exposures

Transgenic lines and wild-type zebrafish were maintained at 28°C, fed brine shrimp twice a day, and kept on regulated by 14/10 hours light/dark cycles. Fish eggs were obtained from natural mating, and embryos were collected and staged based on post-fertilization days. Transgenic zebrafish Tg(*vmat2*:GFP)^36^ expressing green fluorescent protein (GFP) driven by the vesicular monoamine transporter promoter (VMAT2) were used to identify DA, aminergic neurons. Tg(*isl1*[ss]: Gal4-VP16, UAS: EGFP)zf154^37^ transgenic line was used to visualize trigeminal and Rohon-Beard peripheral sensory neurons. To study microglial activation and autophagy, Tg(*mpeg1.1*:mCherry)^38^ and Tg(*elavl3*:eGFP:*map1lc3b*)^39^ lines were utilized respectively. All experiments were conducted following protocols approved by the Animal Research Committee at the University of California, Los Angeles. ZF embryos were manually dechorionated and were exposed to 250nM CPF at 24 hpf at 28°C for 4-6 days.

### γ1-Synuclein knockout zebrafish

Exon 1 of the zebrafish *sncg1* gene encoding γ1-Synuclein ^40^ (also called *sncgb*) was targeted using a pair of custom transcription activator-like effector nucleases (TALENs; supplemental figure [X]A) ^41^. Possible target sequences were found using ZiFiT ^42^ and selected to: (i) encompass a unique restriction site that could be used to identify mutants; and (ii) avoid off-target homology by BLAST search. DNA constructs encoding custom TALENs were generated by iterative assembly using the Joung Lab TAL plasmid set ^41^. The resulting TALEN plasmids were linearized and transcribed *in vitro* to generate mRNA (mMessage Machine T7, Invitrogen). Single cell zebrafish embryos were microinjected with 300 pg mRNA for each TALEN (total 600 pg) in 0.5 – 1.0 nL microinjection buffer ^40^. Surviving embryos were raised to sexual maturity and screened for the germline transmission of *sncg1* deletions by pairwise mating followed by PCR amplification and restriction digest of DNA from progeny embryos. Mating pairs with germline transmission were then outcrossed and their progeny raised to adulthood. F1 founders were identified by fin clip PCR and the mutant allele was cloned and sequenced. We identified 3 different mutant alleles with deletions between 8 – 14bp (supplemental figure [X]A). We selected a 13bp deletion (allele designation Pt131) that is predicted to disrupt the *sncg1* open reading frame (supplemental figure [X]B) for further analysis. The line was expanded from a single F1 founder and outcrossed through four generations prior to analysis to remove any off-target mutations. Heterozygous in-crosses resulted in progeny genotypes at the predicted Mendelian frequency and the *sncg1^Pt131^*mutation was stable over multiple generations. Homozygous *sncg1*^Pt131/Pt131^ zebrafish were viable, fertile and showed no morphological or behavioral deficits during larval development.

Rapid genotyping of the Pt131 allele exploited the deletion of a unique *Afe*I restriction site in exon 1 of sncg1 in the mutant (supplemental figure [X]C). Genomic DNA was extracted from adult fin clips or whole embryos and amplified using primers sncg1F 5’-CCTCTCTCTGTCATTGAAAC-3’ and sncg1R 5’-TAGGAAACACATGCACACAC-3’. The resulting 256bp amplicon was then digested with *Afe*I (New England Biolabs), yielding bands of 164bp and 92bp for the WT allele, and a single band of 243bp for the deletion allele (supplemental figure [X]C).

Western blot analysis of adult brain homogenates showed the expected 14kDa γ1-Synuclein band in WT siblings, but reduced expression in heterozygous *sncg1*^+/Pt131^ zebrafish, and complete loss of γ1-Synuclein expression in homozygous *sncg1*^Pt131/Pt131^ zebrafish (supplemental figure [X]D). These data confirm that Pt131 is a null allele. The homozygous mutant is correspondingly abbreviated to *sncg1*^−/−^ throughout the remainder of the paper.

### ZF behavior assay

ZF larvae were treated from 1 to 7 dpf with 250nM CPF and washed with E3 egg water for 6 hours prior testing. Morphologically normal appearing ZF larvae (7 dpf) were transferred to 96-well plates with 16 larvae per condition, acclimated in the dark for 30 minutes, and their movements were using a ZebraBox (ViewPoint ZebraLab, Civrieux, France). A 10-minute light/10-minute dark cycle was used for 3 cycles for a total of 1 hour of recording. The distances moved by larvae during each increment of 10 minutes were averaged.

### Sample preparation and Liquid Chromatography-Tandem Mass Spectrometry LC-MS

Commercially available CPF was acquired from Chem Service Inc. Diethyl-d10 (CPF-d10) is obtained from CDN Isotopes. Methanol and acetonitrile optima LC/MS grade were purchased from Thermo Fisher Scientific. Standards, controls, and brain tissue samples were prepared in 2 ml beat-beating tubes in duplicates. A five-concentration point (100, 200, 400, 800, 1600 ng/50mg) standard curve and two control samples at low (200 ng/50mg) and high (1600 ng/50mg) concentrations were utilized for compound quantification. Standards, controls, and brain tissue samples stored at -80°C were left to thaw at ambient temperature. To each tube were added six 1.4 mm and three 2.8 mm ceramic beads, 400 ng diethyl-d10 stock solution as an internal standard (IS) prepared in acetonitrile, and 500 mL of a 4:1 mixture acetonitrile/deionized water for extraction. The samples were homogenized in beat-beater for 30 seconds and centrifuged at 16000 xg for five minutes. The supernatant was transferred to new, respectively labeled 1.5 ml microcentrifuges tubes, and samples were dried down in a low-speed vacuum concentrator. The samples were then reconstituted in 100 µl of a methanol/water (v/v 70/30) mixture, vortexed thoroughly, and centrifuged at 16000 x g for five minutes. The supernatant was transferred to HPLC vials. From which 15 µl was injected into the mass spectrometer system for analysis.

Analysis of the analyte concentration was done at the UCLA Pasarow Mass Spectrometry Lab. A targeted LC-MS/MS multiple reaction monitoring (MRM) acquisition method was developed and optimized on a 6460 Agilent Technologies triple quadrupole mass spectrometer. The mass spectrometer was coupled to a 1290 Infinity HPLC system (Agilent Technologies) through a Scherzo SM C-18 analytical column (3 µm 50 x 2 mm UP). For compound elution, a mixture of solvent A (water/formic acid v/v 100/0.1) and solvent B (acetonitrile/formic acid v/v 100/0.1) was used as a mobile phase combined with a linear gradient (min/%B: 0/0, 2/5, 6/100, 7/100, 8/5, 10/5). For the MRM method, the transition from the precursor ion m/z 349.9 to the production m/z 97 isolated for CPF was monitored in positive ion mode at a specific LC retention time. The standard curve was made by plotting the known amount of analyte per standard vs. the ratio of measured chromatographic peak areas corresponding to the analyte over that of the IS (analyte/IS). The trendline equation was used to calculate the absolute concentrations of the analyte in brain tissue.

### Lysotracker labeling and microglial analysis in zebrafish

We used lysotracker staining to image lysosomes in living ZF. Activated microglia have larger and/or more lysosomes than resting microglia. Tg(*mpeg1.1*:mCherry) ZF (5 dpf) which have microglia labeled with mCherry, were incubated in the dark for 45 minutes using 10µM LysoTracker Green DND-26 (ThermoFisher Scientific#L7526) diluted in the treatment solution, and then washed three times in E3 medium before mounting and imaging. Microglial structure was analyzed using FIJI:ImageJ software as previously described.^43^ Briefly, Z-projected images were converted into a 16-bit grayscale file. The images were then made binary, and the skeletonization algorithm at FIJI was used for quantification.

### Measurement of Autophagic Flux

To measure autophagic flux in ZF neurons, we used transgenic [Tg(*elavl3*:eGFP:*map1lc3b*)] ZF larvae as previously described.^39^ Briefly, ZF larvae were treated with 250nM CPF between 24 and 120 hpf. The larvae were anesthetized with 0.01% tricaine and mounted with 1% low melted agarose in a glass bottom culture plate. eGFP-Lc3 punctate were counted with a confocal imaging using a 40x oil immersion objective (numerical aperture = 1.15) and an excitation laser line of 488 nm. Z-stacks comprised 13 sections, and each section of punctate eGFP-Lc3 positive tectum regions was acquired with a slice thickness of 1.5µm and 1024x1024 pixel resolution. For flux assay, zebrafish larvae were incubated with 2µM bafilomycin A1 (Cayman Chemical, 11038) for 55 minutes, and then immediately washed three times with E3 water prior to live imaging. For restoring autophagic flux, larvae were treated with 7.5µM calpeptin (Sigma-Aldrich#C8999) from 48 hpf to 7 dpf with solution refreshed daily.

### Western blot

For analysis of ZF proteins, approximately 40 brains were dissected and homogenized with Radio-Immunoprecipitation Assay (RIPA) lysis buffer with protease inhibitor. After homogenization on ice, samples were centrifuged and protein concentrations in the supernatant were determined using the Bicinchoninic acid (BCA) assay. A final volume of 25 µl was used to load and run 20–30 µg of protein on a 12% SDS-PAGE gel with 1-mercaptoethanol and 1x loading dye. Using XCell-II Blot Module (ThermoFisher Scientific#EI9051), proteins were transferred to the nitrocellulose membrane. Afterward, the transferred membrane was blocked in 5% non-fat milk for 2 hours at room temperature and probed with anti-*Sncg*1 (generated as described previously^44^), anti-Sqstm1/p62 (Cell Signaling Technology#5114; 1:1000 dilution), anti-tubulin (Sigma Aldrich#T9026; 1:2500 dilution) primary antibodies at 4°C overnight. Then the membrane was washed with Tris-buffered saline (50 mM Tris base, 150 mM NaCl, pH 7.5) with Tween-20 (Bio-Rad Laboratories, 1610781) (TBST) followed by donkey-α-rabbit HRP (Santa Cruz Biotechnology#sc-2313; 1:2500 dilution), goat-α-rabbit (Vector Laboratories Burlingame#BA-1000; 1:2500 dilution), and goat-α-mouse HRP (ThermoFisher Scientific#62-6520; 1:2500 dilution) secondary antibodies. Bands were visualized by exposing the blots using chemiluminescent substrate ECL Plus (ThermoFisher Scientific#32132) and then quantified using ImageJ’s gel analysis feature.

### RNA extraction, cDNA preparation, and rtPCR analysis

RNA was extracted from zebrafish larvae at 5 days post fertilization using TRIzol reagent (Invitrogen, 15596026) according to the manufacturer’s instructions and converted into cDNA using iScript™ cDNA Synthesis Kit (Bio-Rad Laboratories#1708890). Gene-specific primer sets (Supplemental Table S1) were used to conduct the real-time (rt) PCR reaction using SsoAdvanced Universal SYBR Green Supermix (Bio-Rad Laboratories#1725270) and cDNA. The 2^-ΔΔcT^ method was used to represent fold change values from cycle number data.

### Statistical analysis

As a part of the PEG analysis, we used univariate, unconditional logistic regression to estimate odds ratios (ORs) and 95% confidence intervals (CIs) for PD and CPF exposure separately by time window and location. As controls, we examined temporal trends in pesticide use based on age, gender, race/ethnicity, study wave, and index year (year of diagnosis or interview). Statistical analyses were performed using Mann-Whitney U-test, and one-way ANOVA with the Bonferroni test where appropriate. Error bars were presented as mean ± standard error of mean (S.E.M). Throughout all studies, a significance level of p < 0.05 was set as significant.

