## Supplementary Figures for "The Pesticide Chlorpyrifos Increases the Risk of Parkinson’s Disease"

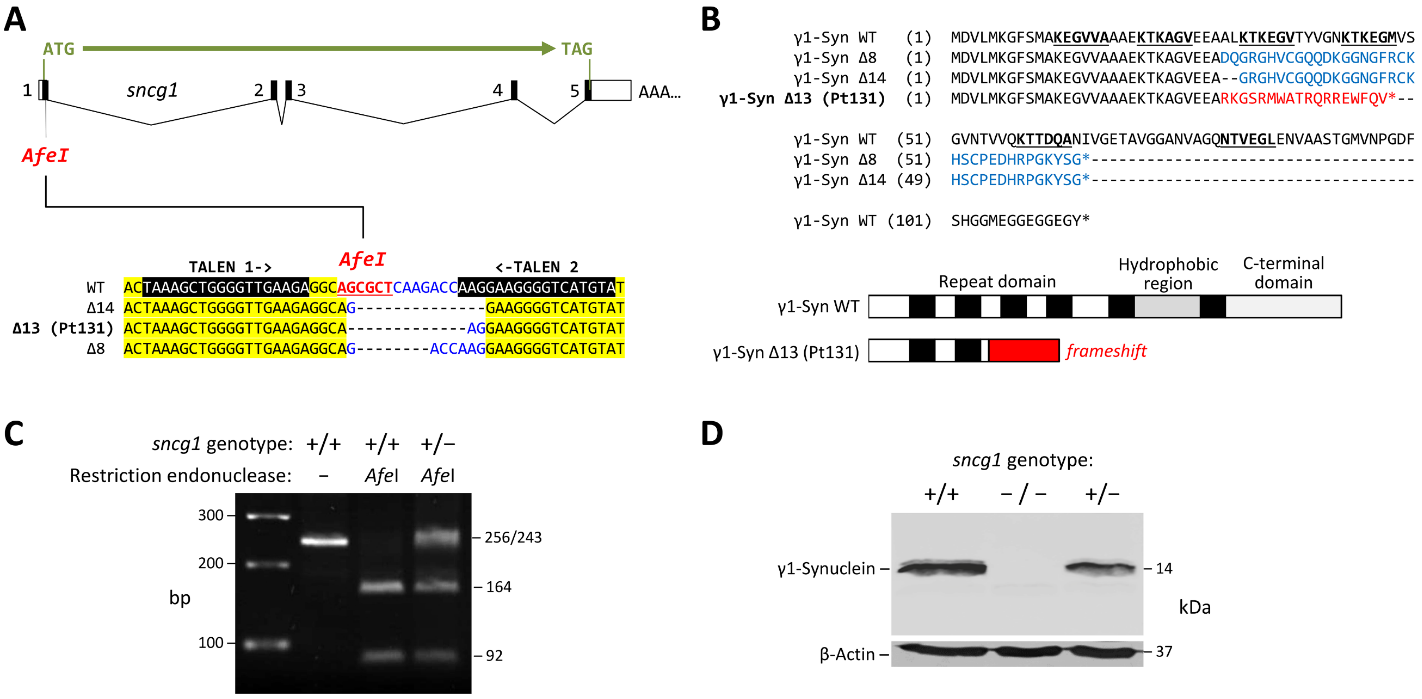


**Supplemental figure 1- γ1-Synuclein knockout zebrafish.** A: γ1-Synuclein knockout zebrafish were generated using TALENs to target an AfeI restriction site in exon 1 of the sncg1 gene. The upper part of the panel shows the genomic structure and open reading frame of the sncg1 gene and mRNA. The sequence alignment below shows the positions and sequences of the two custom TALENs, and the three small deletion alleles found in F1 founders. B: The alignment shows the amino acid sequence of zebrafish γ1-Synuclein with the predicted effects of each of the three deletion alleles shown in panel A. The Synuclein KTKEGV imperfect repeat sequences are underlined, and the frame shift translations of the mutants are shown in blue (frame +1) or red (frame +2). The cartoon below summarizes how the Pt131 mutation truncates γ1-Synuclein near the N-terminus. C: Ethidium-stained 2.5% agarose gel showing PCR products from genomic DNA derived from WT (+/+) or heterozygous (+/−) Pt131 mutants, amplified using *sncg1*-specific primers, and then incubated with *Afe*I, or no enzyme. The WT amplicon of 256bp is cleaved by AfeI into two bands of 164bp and 92bp. The Pt131 mutation removes the *Afe*I site, so there is an uncut band of 243bp in addition to the WT restriction fragments in the heterozygous sample.D: Western blot of adult brain lysates derived from WT(+/+), heterozygous (+/−), or homozygous (−/−) Pt131 mutants. Samples were lysed in RIPA buffer, proteins separated on a 12% polyacrylamide gel and transferred to a nitrocellulose membrane. The upper panel shows the resulting blot probed with a custom polyclonal antiserum to zebrafish γ1-Synuclein, the lower blot shows the same membrane probed with an antibody to β-Actin as a loading control. The Pt131 mutation abolishes expression of γ1-Synuclein, confirming this is a null allele.

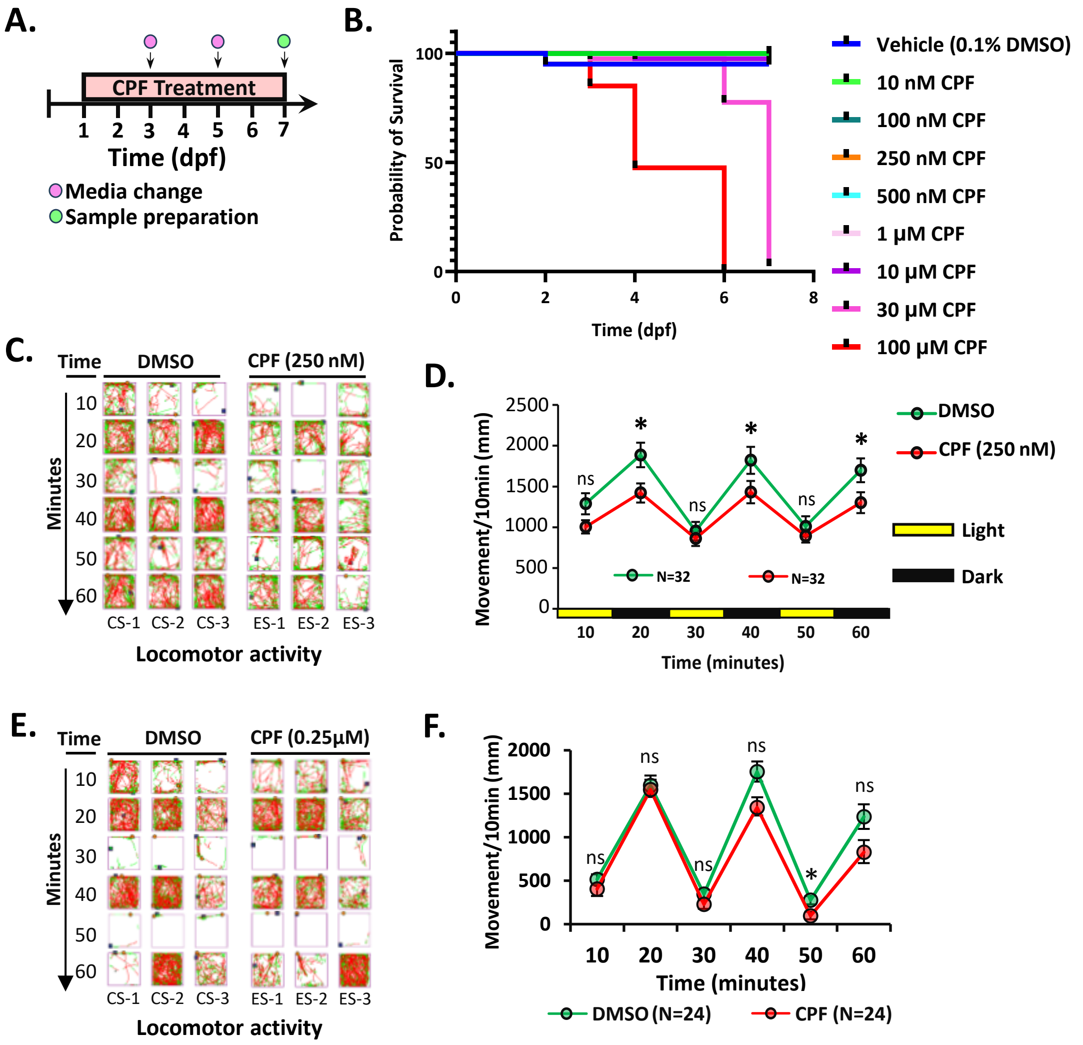


**Supplementary Figure-2: Schematic representation of pesticide treatment and behavior assay in zebrafish embryos.** Tg(*vmat2*:gfp) adult male and female zebrafish were crossed to generate eggs, and at 24 hpf, embryos were treated with 0.25 µM CPF until 7 dpf **(A)**. Embryos were treated with different doses of CPF (10nM to 100µM), and no embryonic lethality was observed until the dose reached 30µM concentration as indicated by the survival curve **(B)**. Motor behavior (distance > 2mm) was tracked for 7dpf zebrafish larvae under an alternating light (yellow)/dark (black) cycle **(C)**. CPF-treated zebrafish larvae traveled significantly less during periods of darkness than vehicle-treated larvae, but there was no significant difference during periods of light **(D)**. **(E, F)** Motor behavior (distance > 2mm) was tracked for 5 dpf zebrafish embryos under an alternating light (yellow)/dark (black) cycle. No significant differences in motor behavior found. Statistical analysis performed by two-tailed, parametric Student t-test. * = $p < 0.05$, ns = not significant. All error bars are presented as the standard error of mean (SEM).

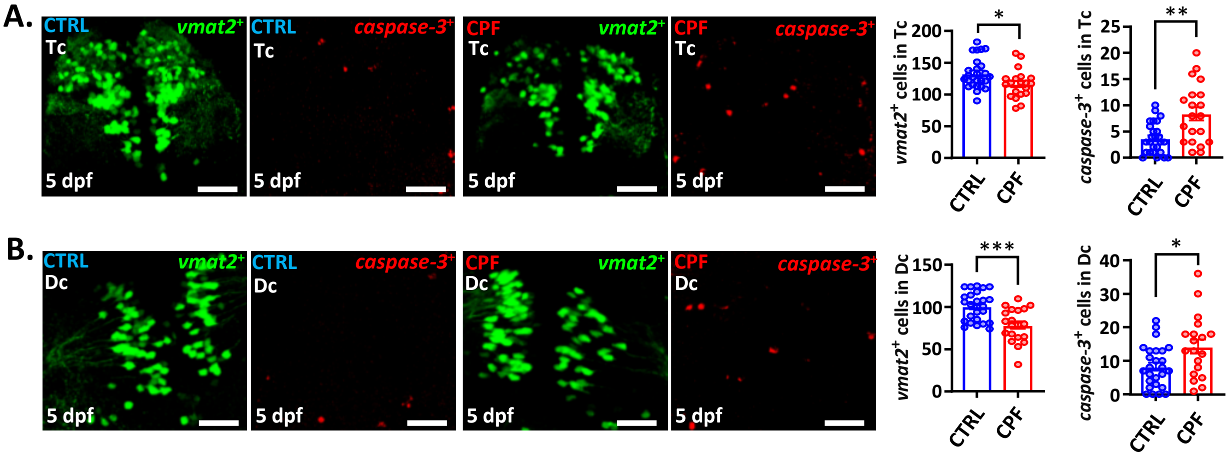


**Supplementary Figure-3: CPF induces apoptosis in zebrafish embryos. (A, B)** Schematic diagram suggested that CPF induces apoptosis in the Tc and Dc region of 5 dpf zebrafish embryos. The number of *caspase-3* positive cells in the midbrain was counted based on Z-Stack (20 layers; each of 2µm thickness) images. Statistical analysis showed the quantification of the numbers of *vmat2*-GFP positive telencephalic and diencephalic clusters of 5 dpf larvae **(B, E).** Scale bar = 50 µm. Statistical analysis performed by two-tailed, non-parametric Mann Whitney U-test. * = $p < 0.05$, ** = $p < 0.01$, *** = $p < 0.001$. All error bars presented as standard error of mean (SEM).

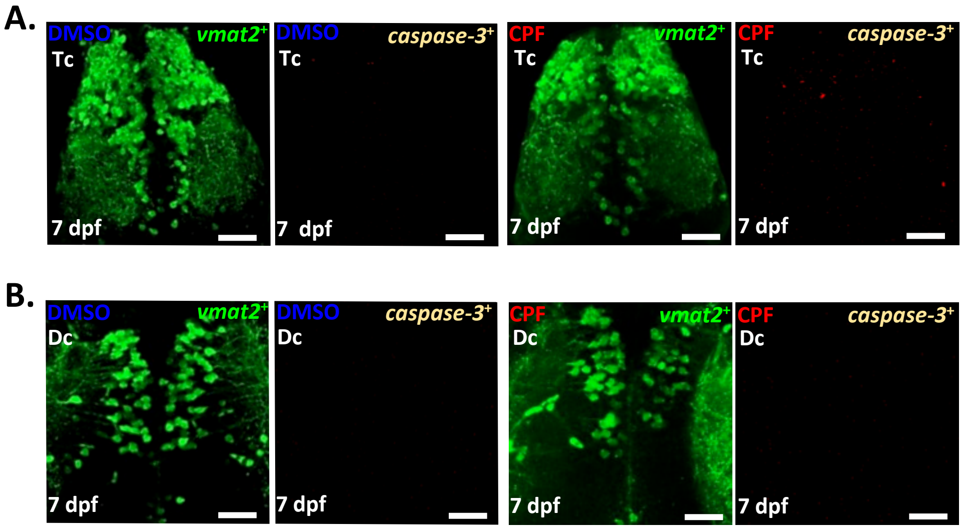


**Supplementary Figure-4: Apoptosis is almost absent in older zebrafish larvae.** Schematic diagram showed no significant apoptosis in the Tc **(A)** and Dc **(B)** region of 7 dpf zebrafish embryos. The number of *caspase-3* positive cells in the telencephalic and diencephalic region was counted based on Z-Stack (20 layers; each of 2µm thickness) images. Scale bar = 50 µm.

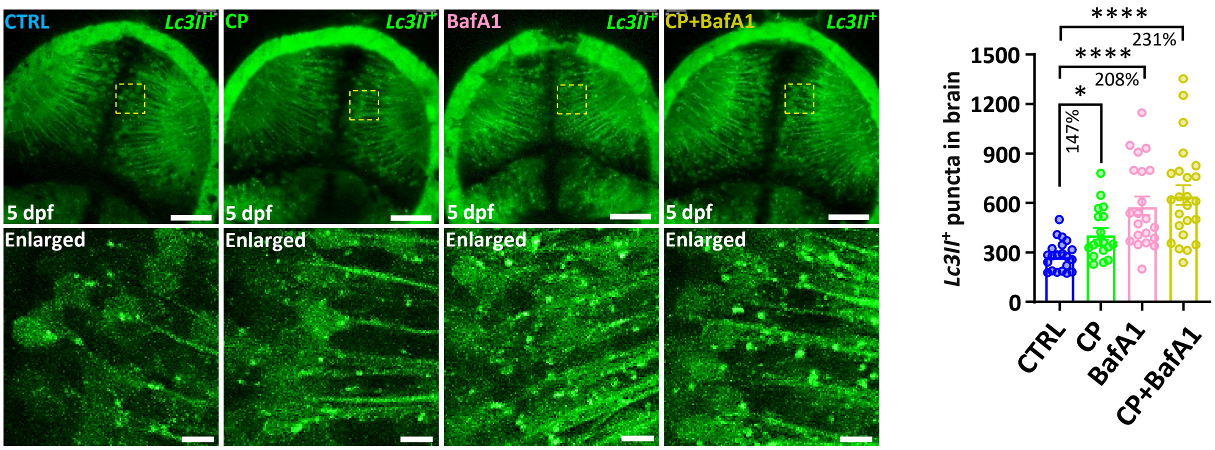


**Supplementary Figure-5: Autophagy induction restores DA neurons in zebrafish embryos.** Schematic diagram showing the position of *Lc3II*-GFP positive cells in the optic tectum regions of 5 dpf larvae. The number of *Lc3II*-GFP positive puncta in the midbrain was counted based on Z-Stack (10 layers out of 15 layers) images. Statistical analysis showed the quantification of the numbers of *Lc3II*-GFP positive puncta in the optic tectum of 5 dpf larvae performed by two-way ANOVA with Brown-Forsythe and Welch's multiple comparisons tests. Scale bar = 50 µm, and 5 µm (Enlarged). CP: Calpeptin. * = $p < 0.05$, **** = $p < 0.0001$, ns = not significant. All error bars are represented as the standard error of mean (SEM).
